# Different components of the RNAi machinery are required for conidiation, ascosporogenesis, virulence, DON production and fungal inhibition by exogenous dsRNA in the Head Blight pathogen *Fusarium graminearum*

**DOI:** 10.1101/633099

**Authors:** Fatima Yousif Gaffar, Jafargholi Imani, Petr Karlovsky, Aline Koch, Karl-Heinz Kogel

**Affiliations:** Department of Phytopathology, Centre for BioSystems, Land Use and Nutrition, Justus Liebig University, Heinrich-Buff-Ring 26, D-35392, Giessen, Germany; Department of Crop Sciences, Molecular Phytopathology and Mycotoxin Research, University of Goettingen, D-37077 Goettingen, Germany

## Abstract

Gene silencing through RNA interference (RNAi) shapes many biological processes in filamentous fungi, including pathogenicity. In this study we explored the requirement of key components of fungal RNAi machinery, including DICER-like 1 and 2 (*Fg*DCL1, *Fg*DCL2), ARGONAUTE 1 and 2 (*Fg*AGO1, *Fg*AGO2), AGO-interacting protein *Fg*QIP (QDE2-interacting protein), RecQ helicase (*Fg*QDE3), and four RNA-dependent RNA polymerases (*Fg*RdRP1, *Fg*RdRP2, *Fg*RdRP3, *Fg*RdRP4), in the ascomycete mycotoxin-producing fungal pathogen *Fusarium graminearum* (*Fg*) for sexual and asexual multiplication, pathogenicity, and its sensitivity to double-stranded (ds)RNA. We corroborate and extend earlier findings that conidiation, ascosporogenesis and Fusarium Head Blight (FHB) symptom development require an operable RNAi machinery. The involvement of RNAi in conidiation is dependent on environmental conditions as it is detectable only under low light (< 2 µmol m^−2^s^−1^). Although both DCLs and AGOs partially share their functions, the sexual ascosporogenesis is mediated primarily by *Fg*DCL1 and *Fg*AGO2, while *Fg*DCL2 and *Fg*AGO1 contribute to asexual conidia formation and germination. *Fg*DCL1 and *Fg*AGO2 also account for pathogenesis as their knock-out (KO) results in reduced FHB development. Apart from KO mutants *Δdcl2* and *Δago1*, mutants *Δrdrp2, Δrdrp3, Δrdrp4, Δqde3* and *Δqip* are strongly compromised for conidiation, while KO mutations in all *RdPRs, QDE3* and *QIP* strongly affect ascosporogenesis. Analysis of trichothecenes mycotoxins in wheat kernels showed that the relative amount of deoxynivalenol (DON), calculated as [DON] per amount of fungal genomic DNA, was reduced in all spikes infected with RNAi mutants, suggesting the possibility that the fungal RNAi pathways affect *Fg*’s DON production in wheat spikes. Moreover, gene silencing by exogenous target gene specific dsRNA (spray-induced gene silencing, SIGS) is dependent on fungal DCLs, AGOs, and QIP, but not on QDE3. Together these data show that in *F. graminearum* different key components of the RNAi machinery are crucial in different steps of fungal development and pathogenicity.

## Introduction

RNA interference (RNAi) is a conserved mechanism triggered by double-stranded (ds)RNA that mediates resistance to exogenous nucleic acids, regulates the expression of protein-coding genes on the transcriptional and post-transcriptional level and preserves genome stability by transposon silencing (Fire et al., 1998; Mello and Conte, 2004; Hammond, 2005; Baulcombe 2013). Many reports have demonstrated that this natural mechanism for sequence-specific gene silencing also holds promise for experimental biology and offers practical applications in functional genomics, therapeutic intervention, and agriculture (Nowara et al., 2010; Koch and Kogel, 2014; Cai et al., 2018; Zanini et al., 2018). Core RNAi pathway components are conserved in eukaryotes, including most parasitic and beneficial fungi (Cogoni and Macino, 1999; Dang et al., 2011; Carreras-Villaseñor et al., 2013; Torres-Martínez and Ruiz-Vázquez, 2017): DICER-like (DCL) enzymes, which belong to the RNase III superfamily, generate double-stranded small interfering (si)RNAs and micro (mi)RNAs (Meng et al., 2017; Song and Rossi, 2017); ARGONAUTE (AGO) superfamily proteins bind small RNA duplexes to form an RNA-induced silencing complex (RISC) for transcriptional and post-transcriptional gene silencing (PTGS) (Zhang et al., 2015; Nguyen et al., 2018); and RNA-dependent RNA polymerases (RdRPs) are involved in the production of dsRNA that initiate the silencing mechanism as well as in the amplification of the silencing signals through the generation of secondary siRNAs (Calo et al., 2012).

Fungal RNAi pathways contribute to genome protection (Meng et al., 2017), pathogenicity (Weiberg et al., 2013; Kusch et al., 2018; Zanini et al. 2019), development (Carreras-Villaseñor et al., 2013), and antiviral defense (Segers et al., 2007; Campo et al., 2016; Wang et al., 2016a). In *Aspergillus flavus* (Bai et al., 2015), *Magnaporthe oryzae* (Raman et al., 2017) and *Penicillium marneffei* (Lau et al., 2013), sRNAs were shown to be responsive to environmental stress. In *Trichoderma atroviride*, both light-dependent asexual reproduction and light-independent hyphal growth require an operational RNAi machinery (Carreras-Villaseñor et al., 2013). Similarly, in *Mucor circinelloides*, defects in the RNAi machinery resulted in various developmental defects such as dysfunction during sexual and asexual reproduction (Torres-Martínez and Ruiz-Vázquez, 2017).

*Neurospora crassa*, a model organism for studying RNAi in filamentous fungi, has different silencing pathways, including quelling (Romano et al., 1992) and meiotic silencing by unpaired DNA (MSUD) (Shiu et al., 2001). In the vegetative stage, the introduction of transgenes results in PTGS of the transgenes and cognate endogenous mRNAs, an RNAi silencing phenomenon known as quelling. The process requires *QDE3* (*Quelling defective 3*), which encodes a RecQ helicase, and RPA (subunit of replication protein A), which recognizes aberrant DNA structures. Interaction of these proteins recruits Quelling defective 1 (QDE1), a protein with dual function as DNA-dependent RNA polymerase (DdRP) and RdPR, to the single-stranded (ss)DNA locus, resulting in production of aberrant ssRNAs and its conversion to dsRNAs. Subsequently the dsRNA is processed into small RNAs by DCL1. sRNAs duplexes are loaded onto QDE2 (Quelling defective 2), which encodes an AGO homolog. QDE2 cleaves the passenger strand and the exonuclease QIP (QDE2-interacting protein) assists to remove it to form an active RISC that targets complementary mRNA for degradation (Chang et al., 2012). MSUD occurs during sexual development in prophase I of meiosis, when unpaired homologous DNA sequences have been detected during the pairing of the homologous chromosomes, which then also leads to the production of aberrant RNA transcripts (Chang et al., 2012). Genes required for MSUD are *SAD1* (*Suppressor of ascus dominance 1*), a paralog of QDE-1, and SAD2. SAD2 recruits SAD1 to the perinuclear region, where aberrant RNA is converted to dsRNA. Upon silencing by DCL1, the small RNA duplexes are loaded onto SMS2 (Suppressor of meiotic silencing 2), an AGO homolog in *Neurospora*, which also is assisted by QIP. In contrast, QDE-2 and DCL2 are not required for MSUD in *Neurospora*, indicating that there are two parallel RNAi pathways functioning separately in the vegetative and meiotic stages.

*Fusarium graminearum* (*Fg*) is one of the devastating pathogens of cereals causing Fusarium Head Blight (FHB) and Crown Rot (FCR) (Dean et al., 2012; Harris et al., 2016). The pathogen belongs to the filamentous ascomycetes. Ascospores are the primary inoculum for FHB epidemics as these spores are forcibly shot into the environment and also can pass long distances (Maldonado-Ramirez et al., 2005). Moreover, the sexual development ensures the formation of survival structures necessary for overwintering (Dill-Macky and Jones, 2000) and the genetic diversity of the population (Cuomo et al. 2007). Of note, spike infections can be symptomless or symptomatic (Urban et al., 2015; Brown et al., 2017). In both cases, *Fusarium* fungi contaminate the grain with mycotoxins and thus decrease grain quality. Among the mycotoxins, the B group trichothecenes, including deoxynivalenol (DON), nivalenol (NIV), and their acetylated derivatives (3A-DON, 15A-DON, and 4A-NIV) influence the virulence of the fungus (Ilgen et al., 2009; Desjardins et al., 1993; Jansen et al., 2005). Mycotoxins such as DON trigger an oxidative burst in the host plants, resulting in cell necrosis and disintegration of the defense system, which then favors colonization of the plant tissues by a necrotrophic fungus (Audenaert et al., 2014). Importantly, *Fg* possesses a functional MSUD mechanism (Son et al., 2011) and *AGO* genes *FgSMS2* or *FgAGO2* are necessary for sexual reproduction (Kim et al., 2015). A recent work discovered that the sex-induced RNAi mechanism has important roles in sexual reproduction (Son et al., 2017). siRNAs produced from exonic gene regions (ex-siRNAs) participate in PTGS at a genome-wide level in the late stages of sexual reproduction. The sex-specific RNAi pathway is primarily governed by *Fg*DCL1 and *Fg*AGO2. Thus, *Fg* primarily utilizes ex-siRNA-mediated RNAi for ascospore formation. Consistent with the key role of *Fg*DCL1 in generative development, the combination of sRNA and transcriptome sequencing predicted 143 novel microRNA-like RNAs (milRNAs) in wild-type perithecia, of which most were depended on *Fg*DCL1. Given that 117 potential target genes were predicted, these perithecium-specific milRNAs may play roles in sexual development (Zeng et al., 2018). To develop RNAi-based plant protection strategies such as host-induced gene silencing (HIGS) (Koch et al. 2013) and spray-induced gene silencing (SIGS) (Koch et al., 2016; Koch et al. 2018) against *Fusarium* species, it is required to bank on knowledge about the RNAi components involved in *Fusarium* development and pathogenicity. A report of Chen and colleagues (Chen et al., 2015) demonstrated that, in *Fg*, a hairpin RNA (hpRNA) can efficiently silence the expression level of a target gene, and that the RNAi components *Fg*DCL2 and *Fg*AGO1 are required for silencing. This finding is consistent with reports showing that a *Fg* wild-type (wt) strain, but not *Fg* RNAi mutants, are amenable to SIGS-mediated target gene silencing, when it grows on a plant sprayed with exogenous dsRNA directed against the fungal *Cytochrome P450 lanosterol C-14α-demethylase* (*CYP51*) genes (Koch et al., 2016). In this study, we expanded previous studies to address the requirement of an extended set of *Fg* RNAi genes in growth, reproduction, virulence, toxin production, and SIGS-mediated inhibition of fungal infection of barley leaves.

## Results

### Requirement of RNAi pathway core components under different light regimes

The *Fg* genome obtained from the Broad Institute (www.broadinstitute.org) contains many functional RNAi machinery components (Chen et al., 2015; Son et al., 2017). We generated *Fg* gene replacement mutants for several major RNAi genes by homolog recombination using the pPK2 binary vector (**Table 1**). Disruption vectors for *FgDCL1, FgDCL2, FgAGO1, FgAGO2, FgRdRP1, FgRdRP2, FgRdRP3, FgRdRP4, FgQDE3*, and *FgQIP* were constructed by inserting two flanking fragments (∼1000 bp) upstream and downstream of the corresponding genes in pPK2 vector (**Table S1**; **Figure S1**). The vectors were introduced into *Agrobacterium tumefaciens*, followed by agro-transformation of the *Fg* strain IFA. Transformants were transferred to Petri dishes of potato extract glucose (PEG) medium, containing 150 μg/ml hygromycin and 150 μg/ml ticarcillin. Mutants were verified by PCR analysis with genomic DNA as template (**Figure 1**) and by expression analysis of the respective RNAi gene (Fig. S2). Colony morphology of PCR verified mutants (12h/12h light/dark, see methods) was inspected in axenic cultures of three different media, PEG, synthetic nutrient (SN) agar and starch agar (SA). In the PEG agar medium, all mutants showed slightly reduced radial growth, while there were no clear differences as compared with the IFA wild type (WT) WT strain in SN and SA media (**Figures S3 A-C**). In liquid PEG medium under day light conditions, all mutants produced comparable amounts of mycelium biomass, though different amounts of the red pigment aurofusarin (Frandsen et al., 2006): *Δdcl1, Δdcl2, Δrdrp1, Δqde3*, and *Δqip1* showed reduced pigmentation, while *Δago1, Δrdrp2, Δrdrp3*, and *Δrdrp4* showed higher pigmentation compared to IFA WT (**Figure S3 D**; **Table 2**). Under light induction conditions (12 h light; 52 µmol m^−2^ s^−1^), conidia grown in 96-well-plate liquid SN cultures showed normal germ tube emergence (not shown). All RNAi mutants formed an elongated hyphal cell type, producing abundant conidia on conidiophores and directly from hyphae. Conidia were moderately curved with clear septations.

**Table 1:** RNAi pathway genes of *Fusarium graminearum* (*Fg*) as identified from www.Broadinstitute.org and used in this study.

| RNAi proteins in *Neurospora crassa* | Homologs in *Fg* | aa identity (%) | Fusarium gene ID | Gene function in *Fg* |
| --- | --- | --- | --- | --- |
| DICER 1 | *FgDCL1* | 43% | FGSG_09025 | Antiviral defence (Wang et al., 2016a).<br>Minor role in processing of exogenous dsRNA, hpRNA or pre-miRNA in mycelium (Chen et al., 2015).<br>Major role in sex-specific RNAi pathway: Production of regulatory sRNAs. Required for ascospore production (Son et al., 2017). |
| DICER 2 | *FgDCL2* | 35% | FGSG_04408 | Processing of exogenous dsRNA, hpRNA and pre-miRNA in mycelium (Chen et al., 2015).<br>Partially shared DCL-1 role in production of regulatory sRNAs in the sexual stage (Son et al., 2017). |
| ARGONAUTE 1<br>(syn. Quelling defective 2) | *FgAGO1* | 59% | FGSG_08752 | Major component in the RISC during quelling (Chen et al., 2015). |
| ARGONAUTE 2<br>(syn. Suppressor of meiotic silencing 2, SMS2) | *FgAGO2* | 43% | FGSG_00348 | Minor role in binding siRNA derived from exogenous dsRNA, hpRNA or pre-miRNA in mycelium (Chen et al., 2015).<br>Major role in sex-specific RNAi pathway; required for ascospore production (Son et al., 2017). |
| RNA-DEPENDENT<br>RNA POLYMERASE<br>(syn. Quelling defective 1) | *FgRdRP1* | 38% | FGSG_06504 | Maybe associated with secondary sRNA production (Chen et al., 2015). |
|  | *FgRdRP4* | 33% | FGSG_04619 | Maybe associated with secondary sRNA production (Chen et al., 2015). |
| RNA-DEPENDENT<br>RNA POLYMERASE<br>(syn. Suppressor of ascus dominance, SAD1) | *FgRdRp2* | 42% | FGSG_08716 | Maybe associated with secondary sRNA production (Chen et al., 2015). |
|  | *FgRdRp5* | 29 % | FGSG_09076 | Roles in the antiviral defence.<br>Maybe associated with secondary sRNA production (Chen et al., 2015). |
| RNA-DEPENDENT<br>RNA POLYMERASE<br>(RRP3) | *FgRdRP3* | 47% | FGSG_01582 | Maybe associated with secondary sRNA production (Chen et al., 2015). |
| QDE2-INTERACTING<br>PROTEIN | *FgQIP* | 32% | FGSG_06722 | The homolog has been identified in (Chen et al., 2015), but not yet studied in depth. |
| RecQ HELICASE<br>QDE3 | *FgQDE3* | 46% | FGSG_00551 | not studied. |

**Table 2:**
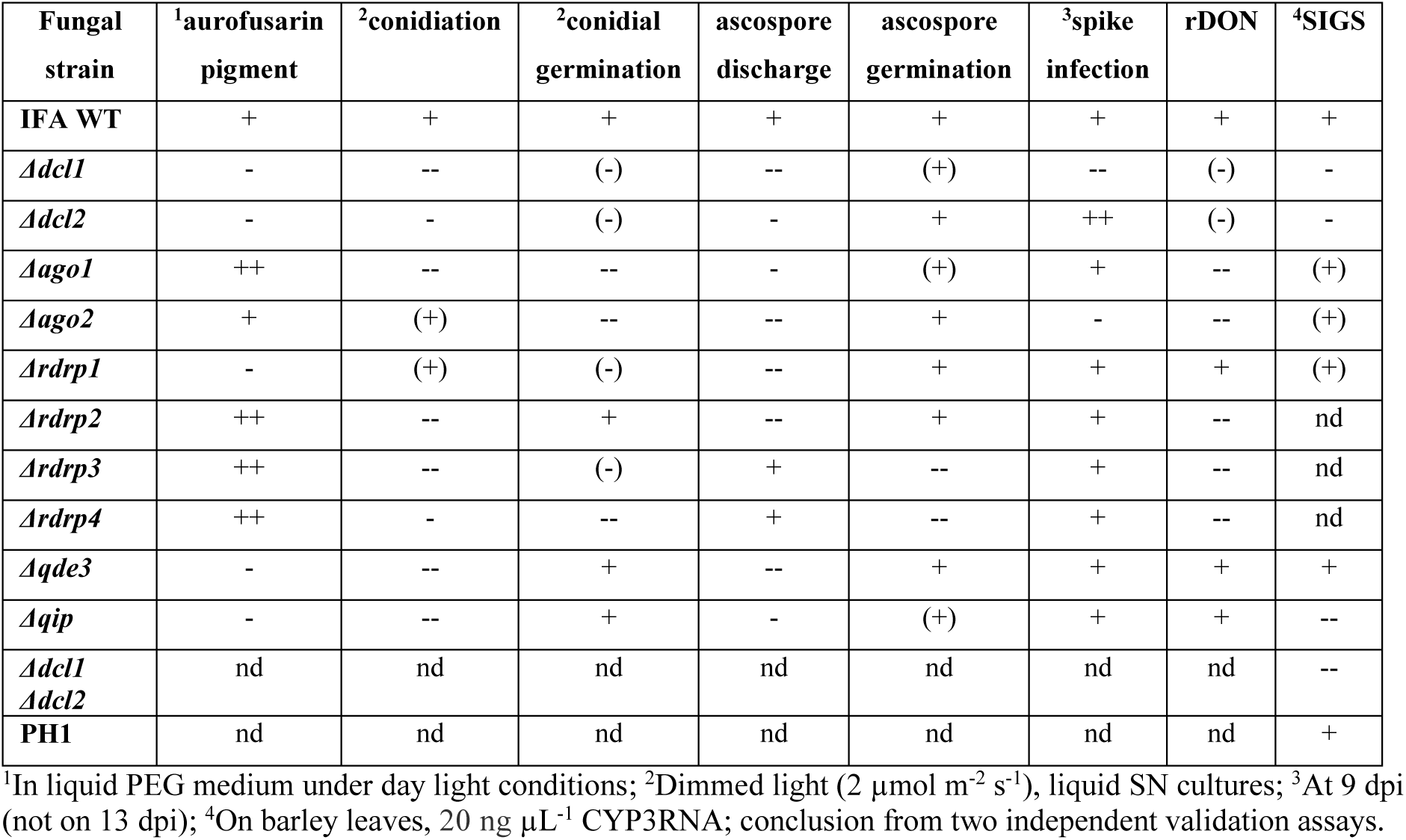
Function of *Fusarium graminearum* RNAi mutants in various developmental and pathogenic processes.

**Fig. 1.**
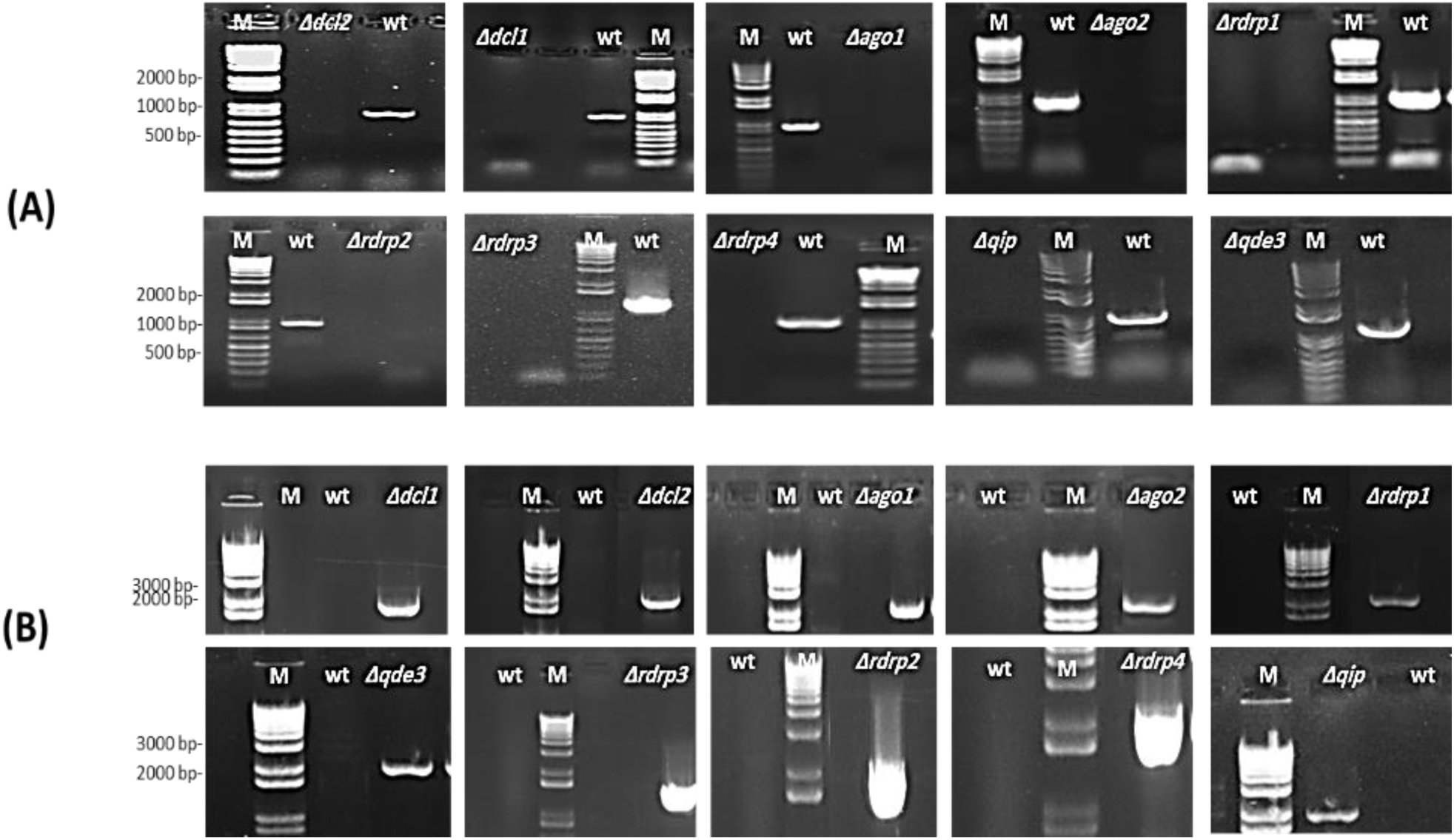
PCR verification of targeted gene replacement in *Fusarium graminearum*. **(A)** Amplification of an internal part of the targeted genes *DCL1, DCL2, AGO1, AGO2, RdRP1, RdRP4, RdRP2, RdRP3, QIP*, and *QDE3* are positive in IFA WT and negative in corresponding mutants. (**B)** PCR with primer pairs in the right recombination sequence and hygromycin, showing that the antibiotic resistance gene had integrated into the target gene locus. PCR products were analyzed on 1.5% agarose gel electrophoresis. M; DNA marker. wt; wild type.

When grown continuously under dimmed light (2 µmol m^−2^ s^−1^), liquid SN cultures of RNAi mutants showed significantly reduced conidiation compared to IFA WT, except *Δago2* and *Δrdrp1*, which were only slightly affected (**Figure 2 A**). Under this non-inductive condition, some RNAi mutants also were compromised in conidial germination: *Δago1, Δago2* and *Δrdrp4* showed significantly reduced germination, while *Δrdrp3, Δdcl1, Δrdrp1* and *Δdcl2* showed a slight reduction, and *rdrp2, Δqip* and *Δqde3* showed normal conidial germination (**Figure 2 B**; see **Table 2**). All RNAi mutants had a normal germ tube morphology, except *Δrdrp4*, which tends to develop multiple germ tubes (**Figure 2 C**). These results suggest a requirement for *Fg* RNAi components genes in the full control of asexual development depending on the environmental conditions.

**Fig. 2.**
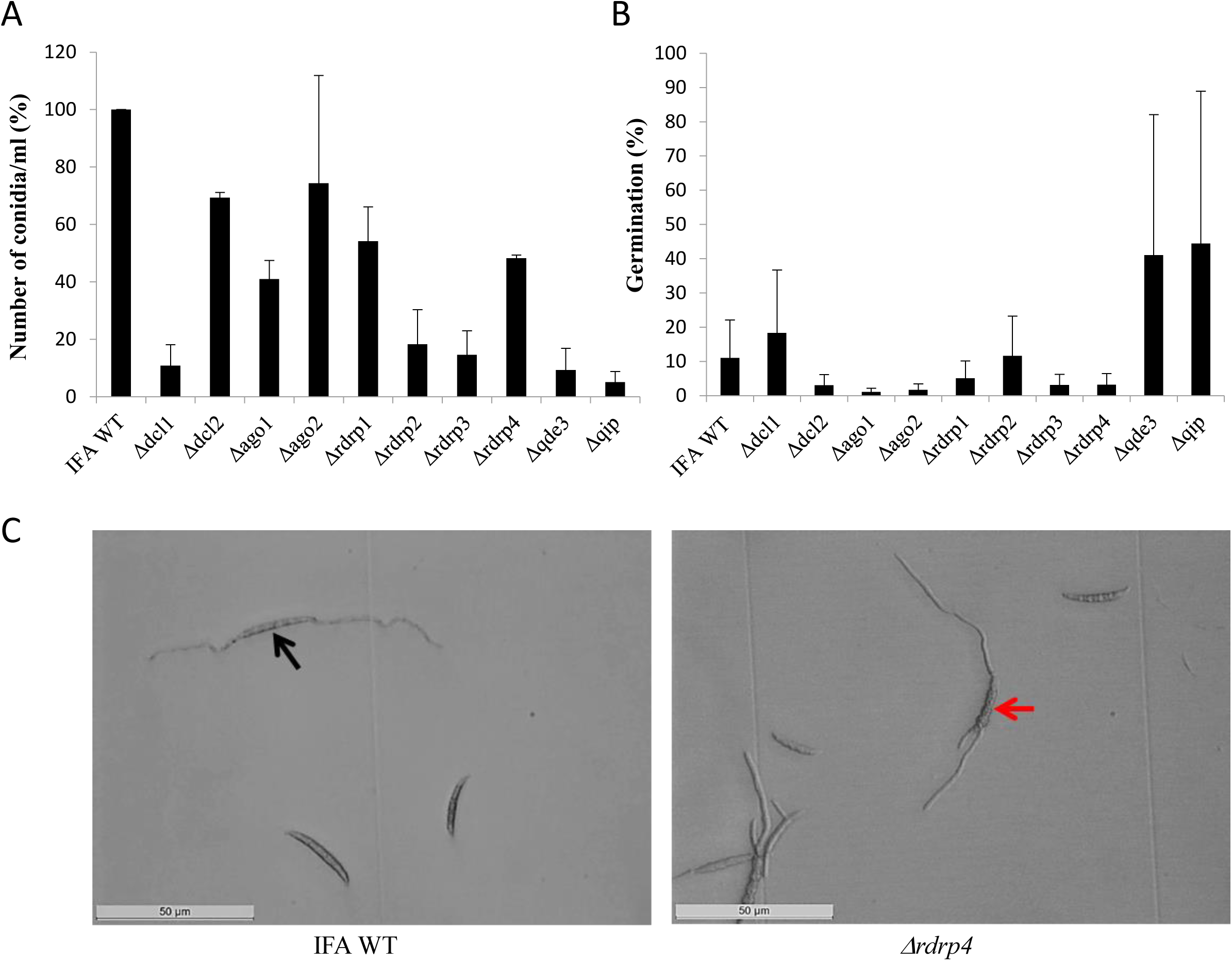
The RNAi pathway is required for asexual development of *Fusarium graminearum* in the absence of inductive light. **(A)** Number of conidia produced: Means± SEs of the percentage of conidia numbers from three repeated experiments. Significant differences are marked: *P, 0.05, **P, 0.01, ***P, 0.001 (Student’s *t* test). **(B)** Percent of conidial germination: Means± SEs of the percentage of germinated spores from three biological repetitions. Significant differences are marked: *P, 0.05 (Student’s t test). **(C)** Microscopic observation of germinated and non-germinated conidia of IFA WT and *Δrdrp4*. Imaging after 48 h incubation in the dark, scale bar: 50 μm. Black arrow; conidia forming a bipolar germ tube. Red arrow; conidia forming multiple germ tubes.

### *F. graminearum* RNAi components are required for sexual development

Because there were contrasting data in the literature, we resumed asking the question of whether RNAi components are required for sexual reproduction of *Fg*. Perithecia (fruiting bodies) formation was induced in axenic cultures on carrot agar (Cavinder et al., 2012). All RNAi mutants produced melanized mature perithecia to the same extend as compared to IFA WT (not shown). Next, we assessed the forcible discharge of ascospores by a spore discharge assay (**Figure 3**). Discharge of ascospores from perithecia into the environment results from turgor pressure within the asci; the dispersal of ascospores by forcible discharge is a proxy for fungal fitness as it is important for dissemination of the disease. To this end, half circular agar blocks covered with mature perithecia were placed on glass slides and images from forcibly fired ascospores (white cloudy) were taken after 48 h incubation in boxes under high humidity and fluorescent light. We found that the forcible discharge of ascospores was severely compromised in *Δdcl1, Δago2, Δrdrp1, Δrdrp2, Δqde3*, and less severe in *Δdcl2, Δago1, Δqip1*, while *Δrdrp3* and *Δrdrp4* were indistinguishable from IFA WT. (**Figures 3 A, B**). Microscopic observation of the discharged ascospores revealed that their morphology was not affected (not shown). However, the percentage of discharged ascospores that retained the ability to germinate varied in the mutants with *Δrdrp3* and *Δrdrp4*, showing strong reduction in the ascospore germination (**Figure 3 C**; see **Table 2**). Together, these results confirm that the RNAi pathway is involved in sexual reproduction, though the requirement of individual RNAi components greatly varies in the different developmental stages.

**Fig. 3.**
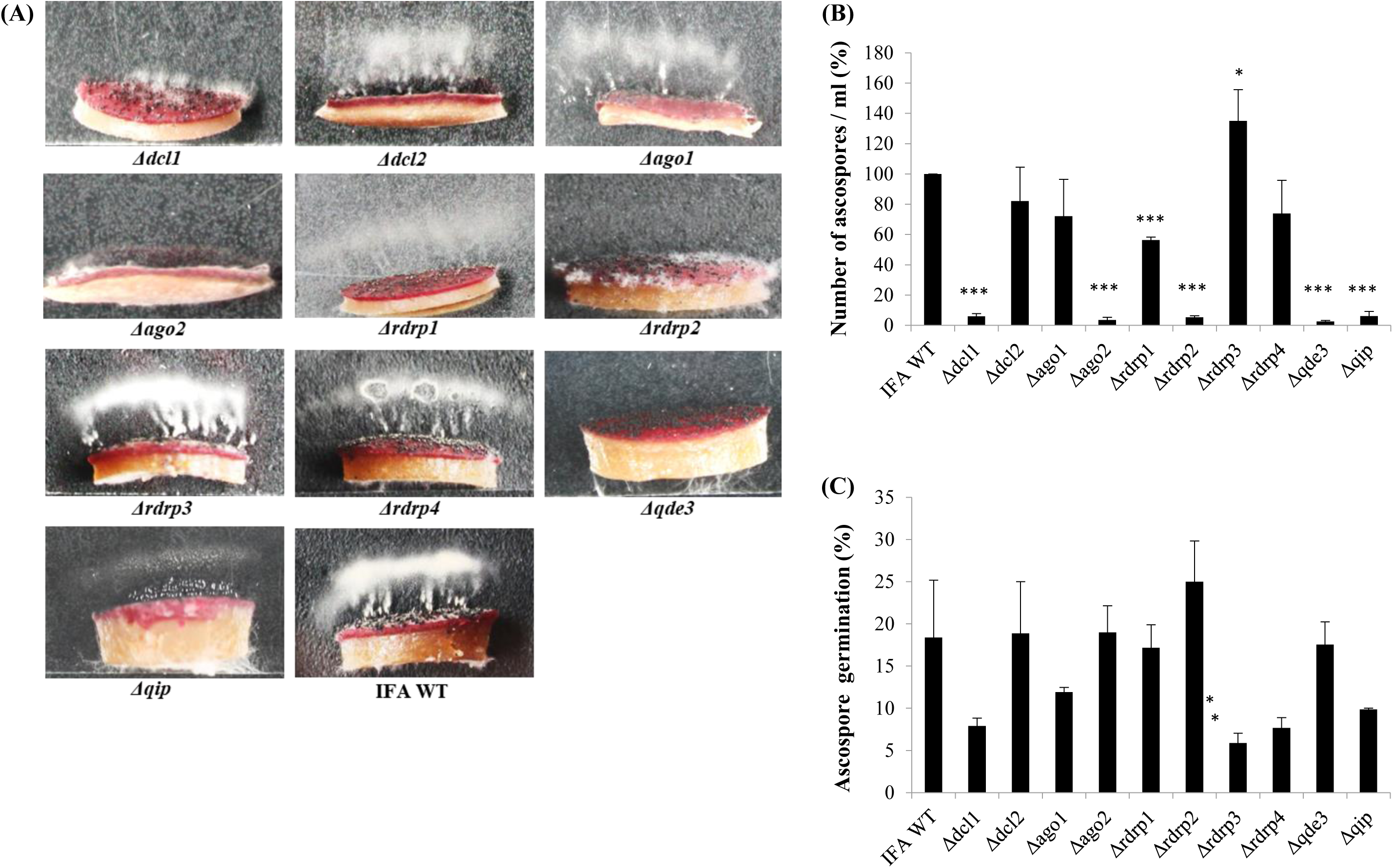
Forcible ascospore discharge in *Fusarium graminearum* RNAi mutants and IFA WT. **(A)** Forcible ascospore firing. Half circular carrot agar blocks covered with mature perithecia were placed on glass slides. Photos from forcibly fired ascospores (white cloudy) were taken after 48 h incubation in boxes under high humidity and fluorescent light. **(B)** Fired ascospores were washed off and counted. Means± SDs of the counted spores is presented from three biological repetitions. Significant differences are marked: *P, 0.05, ***P, 0.001 (Student’s t test). **(C)** Ascospore germination. Discharged ascospores were incubated at 100% RT in the dark for 24 h at 23°C in SN liquid medium. The percentage of germination was assessed by examining the ascospore number in three random squares in the counting chamber. Means± SEs of the percentage of germinated spores from three biological repetitions. Significant differences are marked: *P, 0.05 (Student’s t test).

### *F. graminearum* RNAi mutants show variation in kernel infection

It has been reported that *Fg* mutants defective in DCL, AGO, or RdRP were not compromised in virulence on wheat spikes (Chen et al., 2015). We extended this previous study by testing additional *Fg* RNAi mutants. Conidia were point-inoculated to a single spikelet at the bottom of a spike of the susceptible wheat cultivar Apogee. Fungal colonization was quantified nine and 13 days post inoculation (dpi) by determining the infection strength. Infected parts of a spike bleached out, whereas the non-inoculated spikes remained greenish. At late infection stages (13 dpi), all RNAi mutants caused strong FHB symptoms comparable with IFA WT. However, we found clear differences in the severity of infections at earlier time points (9 dpi), with *Δdcl1* and *Δago2* showing most compromised FHB development (**Figure** 4 A; see **Table 2**). At 13 dpi, RNAi mutants also showed considerable variation on *Fg-*infected kernel morphology (**Figure S4 A**). Thousand-grain-weight (TGW) of kernels infected with RNAi mutants showed slight, though not significant differences, in the total weights compared to IFA WT infection (**Figure S4 B**).

**Fig. 4.**
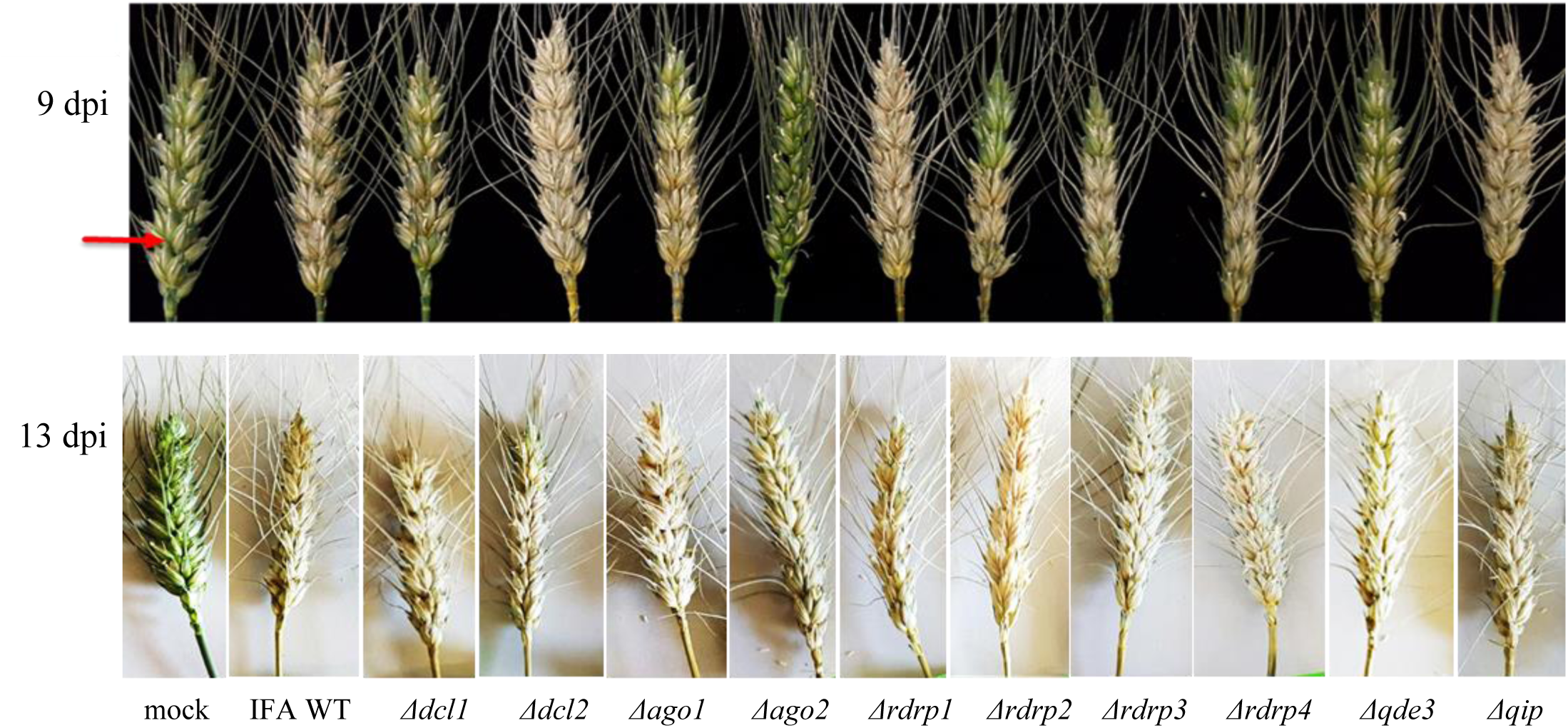
Infection of Apogee wheat spikes with *Fusarium graminearum* RNAi mutants and IFA WT. **(A)** Representative samples of spikes at 9 and 13 dpi. One spikelet at the bottom of each spike (red arrow) was point inoculated with 5 µl of 0.002% Tween 20 water containing40,000 conidia / mL. The assay was repeated two times with 10 spikes per fungal genotype and experiment. **(B)** Wheat kernels 13 dpi with *Fg* RNAi mutants and IFA WT.

### DON production is compromised in *F. graminearum* mutants that show reduced pathogenicity on wheat kernels

We quantified the amount of DON in *Fg*-infected wheat spikes at 13 dpi (point-inoculation using 5 µl of 0.002% Tween 20 water containing 40,000 conidia / mL) at mid-anthesis. Of note, the relative amount of DON [rDON], calculated as [DON] relative to the amount of fungal genomic DNA, was reduced in virtually all spikes infected with RNAi mutants, whereby spikes infected by *Δqip* and *Δdcl2* showed the lowest toxin reduction as compared with the other mutants (**Table 3**). The data suggest that fungal RNAi pathways affect *Fg*’s DON production in wheat spikes. While [rDON] changed, the ratio of [DON] and [A-DON] (comprising 3A-DON and 15A-DON) remained constant in all mutants vs. IFA WT, suggesting that the fungal RNAi pathways do not affect the trichothecene metabolism.

**Table 3:** Trichothecenes produced by RNAi mutants in infected wheat kernels at 13 dpi.

| Samples | ng *Fg* DNA /mg seed d.w. | DON [mg/kg seed] | DON/DNA | <sup>1</sup>A-DON [mg/kg seed] | <sup>2</sup>A-DON/DON x1000 |
| --- | --- | --- | --- | --- | --- |
| Mock (without *Fg*) | 0 | 0.00 | 0 | 0 | 0 |
| *Δago1* | 0.84 | 12.7 | 15.2 | 0.45 | 36 |
| *Δago2* | 1.28 | 32.3 | 25.2 | 1.04 | 32 |
| *Δdcl1* | 2.86 | 61.9 | 21.6 | 1.87 | 30 |
| *Δdcl2* | 2.03 | 56.6 | 27.9 | 1.89 | 33 |
| *Δrdp1* | 4.84 | 86.7 | 17.9 | 3.51 | 40 |
| *Δrdp2* | 0.95 | 16.9 | 17.8 | 0.43 | 25 |
| *Δrdp3* | 0.78 | 12.3 | 15.6 | 0.35 | 29 |
| *Δrdp4* | 0.47 | 4.90 | 10.3 | 0.15 | 31 |
| *Δqip* | 2.53 | 68.7 | 27.2 | 2.87 | 42 |
| *Δqde3* | 4.33 | 82.3 | 19.0 | 3.91 | 47 |
| IFA WT | 2.18 | 78.3 | 35.9 | 2.58 | 33 |

DON, deoxynivalenol; A-DON, acetyldeoxynivalenol.
<sup>1</sup> 3A-DON (3-acetyldeoxynivalenol) and 15A-DON (15-acetyldeoxynivalenol) were measured
<sup>2</sup> Ratio of concentrations of A-DON and DON, multiplied by 1000

### *F. graminearum* RecQ helicase mutant *Δqde3* is insensitive to dsRNA

Spraying plant leaves or fruits with dsRNA targeting essential fungal genes can reduced fungal infections by spray-induced gene silencing (SIGS) (Koch et al., 2016; Wang et al., 2016b; Dalakouras et al., 2016; McLoughlin et al., 2018). We addressed the question which RNAi mutants are compromised in SIGS upon treatment with dsRNA. To this end, we conducted a SIGS experiment on detached barley leaves that were sprayed with 20 ng µL^−1^ CYP3RNA, a 791 nt long dsRNA that targets the three fungal genes *FgCYP51A, FgCYP51B* and *FgCYP51C* (Koch et al., 2016). By 48 h after spraying, leaves were drop inoculated with 5 × 10^4^ conidia ml^−1^ of *Fg* RNAi mutants and IFA WT. Five days later, infected leaves were scored for disease symptoms and harvested to measure the expression of the fungal target genes by qPCR (**Figure 5**). As revealed by reduced disease symptoms, leaves sprayed with CYP3RNA vs. TE (buffer control), only *Δqde3* was equally sensitive to dsRNA like the IFA WT, while all other mutants tested in this experiment were slightly or strongly compromised in SIGS and less sensitive to CYP3RNA (**Figure 5 A**, see **Table 2**). Consistent with this, strong down-regulation of all three *CYP51* target genes was observed only in IFA WT and *Δqde3.* In *Δdcl1, Δdcl2* and *Δqip1*, the inhibitory effect of CYP3RNA on *FgCYP51A, FgCYP51B* and *FgCYP51C* expression was completely abolished (**Figure 5B**). To further substantiate this finding, we tested a *dcl1/dcl2* double mutant in *Fg* strain PH1. As anticipated from the experiments with IFA WT, the PH1 *dcl1/dcl2* mutant was fully compromised in SIGS (**Figures 5A, B**).

**Fig. 5.**
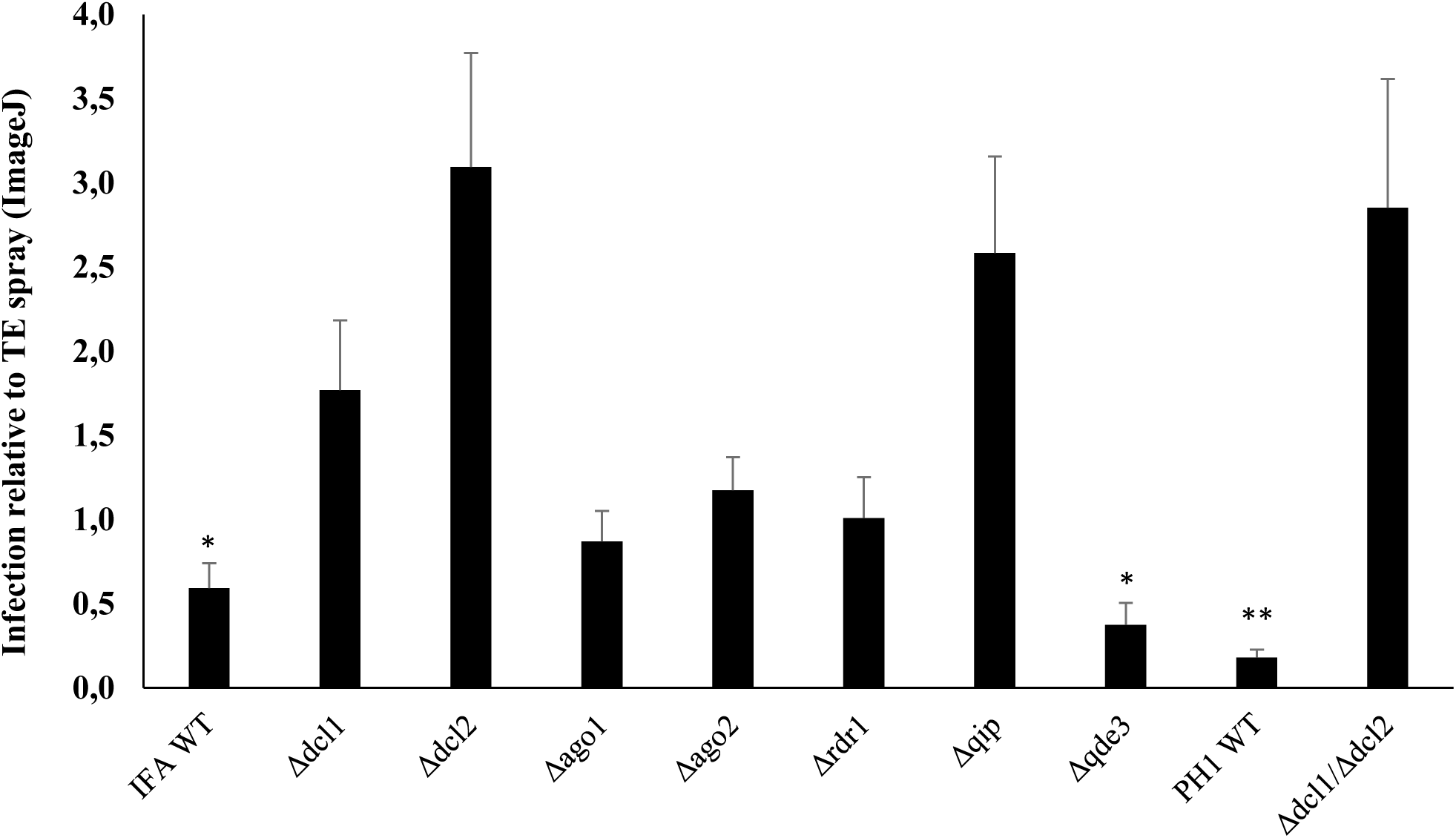
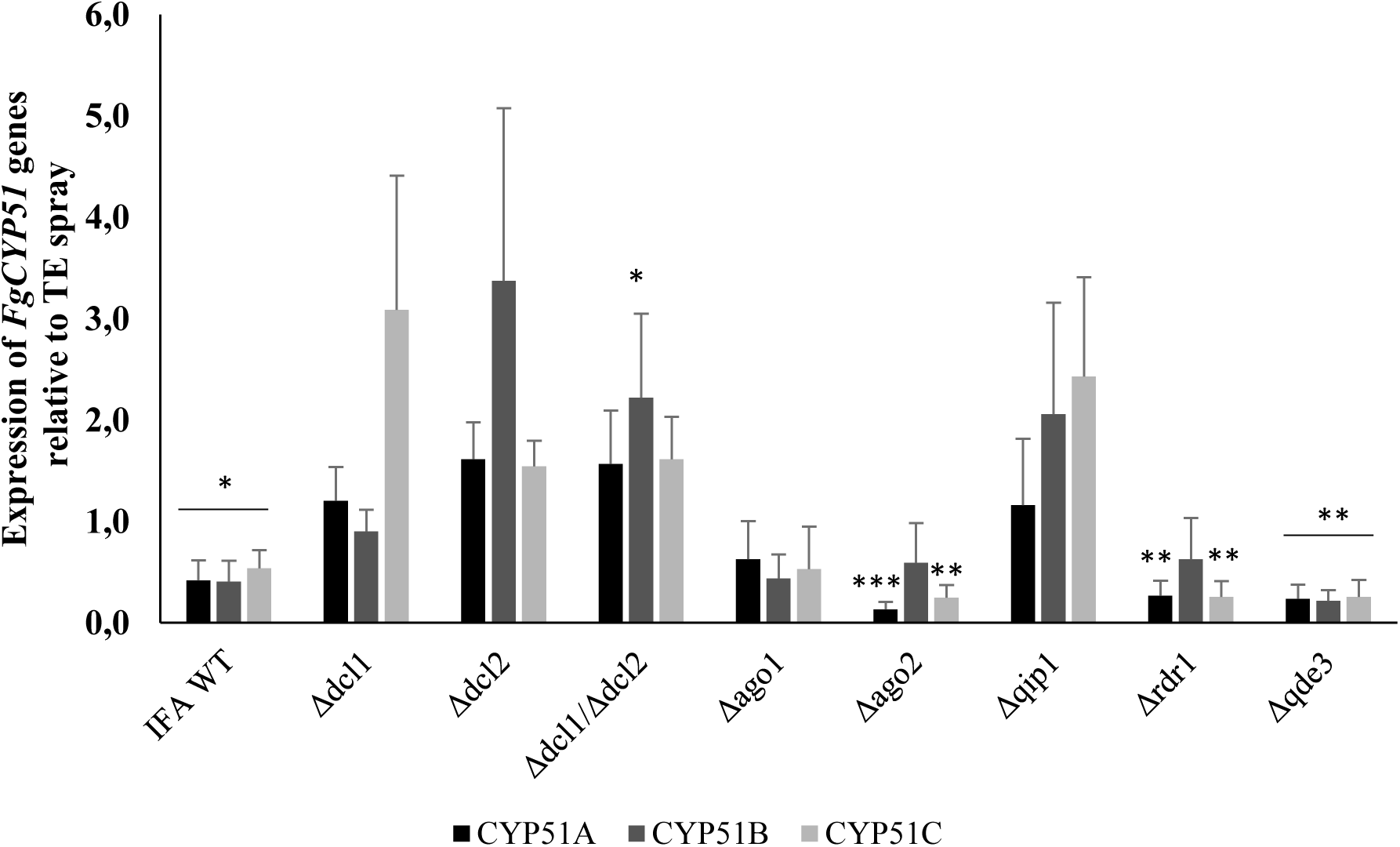
Infection symptoms of *Fg* RNAi mutants on barley leaves sprayed with the dsRNA CYP3RNA. **A.** Detached leaves of three-week-old barley plants were sprayed with 20 ng μl^−1^ CYP3RNA or TE buffer, respectively. After 48 h, leaves were drop-inoculated with 5 × 10^4^ conidia ml^−1^ of indicated *Fg* RNAi mutants and evaluated for infection symptoms at 5 dpi. Values show relative infection area as calculated from TE-vs. CYP3RNA-treated plants for each RNAi mutant with 10 leaves and thee biological repetitions. Asterisks indicate statistical significant reduction of the infection area on CYP3RNA-vs. TE-treated plants measured by ImageJ for each mutant (**p<0,01; ***p< 0,001; students t-test). The *dcl1 dcl2* double mutant is generated in *Fg* strain PH1. (**B**). Downregulation of the three *CYP51* genes in *Fg* mutants upon colonization of CYP3RNA-vs. TE-treated barley leaves. Asterisks indicate statistical significant downregulation of *CYP51* genes on CYP3RNA vs. TE-treated plants. (**p<0,01; ***p< 0,001; students t-test). Error bars indicate SE of three independent experiments in A and B.

## Discussion

We generated a broad collection of knock-out mutants for RNAi genes in the necrotrophic, mycotoxin-producing pathogen *Fusarium graminearum* to demonstrate their involvement in vegetative and generative growth, disease development, mycotoxin production, and sensitivity to environmental RNAi. A summary of the mutants’ performance in the various processes is shown in Table 2. While all RNAi mutants show normal vegetative development in axenic cultures, there were differences in pigments production in liquid potato extract glucose cultures. This suggests that in *Fg* an RNAi pathway regulates the gene cluster responsible for the biosynthesis of pigments, including aurofusarin. Aurofusarin is a secondary metabolite belonging to the naphthoquinone group of polyketides that shows antibiotic properties against filamentous fungi and yeast (Medentsev et al., 1993). The function of the compound in the fungus is unknown as white mutants have a higher growth rate than the wt and are as pathogenic on wheat and barley (Malz et al., 2005).

Overall, the contribution of the RNAi pathways to vegetative fungal development and conidiation varies among different fungi and must be considered case by case. Under low light (< 2 µmol m^−2^ s^−1^) all *Fg* RNAi mutants showed reduced conidia production and some showed aberrant germination compared to IFA WT. This suggests that in the absence of light induction the RNAi pathway is required for conidiation. RNAi may play a role in regulation of light responsive genes affecting conidiation as shown for *T. atroviride*, where DCL2 and RdRP3 control conidia production under light induction (Carreras-Villaseñor et al., 2013). The authors claimed that *Δdcl2* and *Δrdrp3* are impaired in perception and/or transduction of the light signal affecting the transcriptional response of light-responsive genes. Similarly, *Metarhizium robertsii dcl* and *ago* mutants show reduced abilities to produce conidia under light, though the light quantity was not described (Meng et al., 2017).

Perithecia development has been used to study sexual development and transcription of genes related to sexual development (Trail et al., 2000; Qi et al., 2006; Hallen et al., 2007). In field situations, ascospores serve as the primary inoculum for FHB epidemics because these spores are shot into the environment and can spread long distances (Maldonado-Ramirez et al., 2005). We found that all RNAi mutants could produce mature perithecia. However, corroborating and extending the exemplary work of Son et al. (2017), we also found that, beside *FgDCL1* and *FgAGO2*, other RNAi genes such as *RdRP1, RdRP2, RdRP3, RdRP4, QDE3, QIP* contribute to the sexual reproduction. Mutations in these genes either showed severe defect in forcible ascospore discharge or significantly reduced germination. The Son et al. (2017) study showed that *Fgdcl1* and *Fgago2* are severely defective in forcible ascospore discharge, while *Fgdcl2* and *Fgago1* show indistinguishable phenotypes compared to the wt. Active roles for *Fg*DCL1 and *Fg*AGO2 is supported by the finding that expression levels of many genes, including those closely related to the mating-type (*MAT*)-mediated regulatory mechanism during the late stages of sexual development, was compromised in the respective mutants after sexual induction (Kim et al., 2015). Moreover, *Fg*DCL1 and *Fg*AGO2 participate in the biogenesis of sRNAs and perithecia-specific miRNA-like RNAs (milRNAs) also are dependent on *Fg*DCL1 (Zeng et al., 2018). Most of the produced sRNA originated from gene transcript regions and affected expression of the corresponding genes at a post-transcriptional level (Son et al., 2017).

While our data show that, in addition to *Fg*DCL1 and *Fg*AGO2, more *Fg* RNAi-related proteins are required for sex-specific RNAi, further transcriptomic analysis and sRNA characterization are needed for a mechanistic explanation. Of note, ex-siRNA functions are important for various developmental stages and stress responses in the fungus *M. circinelloides*, while *Fg* utilizes ex-siRNAs for a specific developmental stage. Thus, ex-siRNA-mediated RNAi might occur in various fungal developmental stages and stress responses depending on the fungal species.

We investigated the involvement of RNAi in pathogenicity and FHB development by infecting wheat spikes of the susceptible cultivar Apogee with fungal conidia. At earlier time points of infection (9 dpi) clear differences in virulence between RNAi mutants were observed, though all mutants could spread within a spike and caused typical FHB symptoms at later time points (13 dpi). Despite full FHB symptom development in all mutants at 13 dpi, we observed various effects of fungal infection on the kernel morphology, corresponding to the different aggressiveness of mutants at early time points. Since this phenomenon may account for differences in producing mycotoxins during infection, we quantified mycotoxins in the kernels. Of note, [rDON] was reduced in virtually all spikes infected with RNAi mutants, whereby [rDON] was strongly reduced especially in spikes colonized with mutants *Δago1, Δrdrp1, Δrdrp2, Δrdrp3, Δrdrp4* and *Δqde3* as compared with IFA WT (see Table 2 and Table 3). The data suggest that fungal RNAi pathways affect *Fg*’s DON production in wheat spikes. While [rDON] changed, the ratio of [DON] and [3A-DON] remained constant in all mutants vs. IFA WT, suggesting that the fungal RNAi pathways do not affect the trichothecene chemotype.

Our work also identifies additional *Fg* RNAi proteins associated with sensitivity to dsRNA treatments. For the validation of the effects we used two independent tests: infecting phenotyping and qRT-PCR analysis of fungal target genes. *Δdcl1* and *Δdcl2* as well as *Δqip* and *Δrdrp* showed compromised SIGS phenotypes in either tests, strongly suggesting that these proteins are required for environmental RNAi in *Fg*. Further substantiating our finding, the fungal *dcl1/dcl2* double mutant of *Fg* strain PH1 also showed complete insensitivity to dsRNA and thus is fully compromised to environmental RNAi.

Taken together, our results further substantiate the involvement of RNAi pathways in conidiation, ascosporogenesis and pathogenicity of *Fg*. Nevertheless, further studies must explore the mechanistic roles of *Fg* RNAi genes in these processes.

## Methods

### Fungal material, generation of gene deletion mutants in *Fusarium graminearum*

The *Fg* strain PH1 and the PH1 *dcl1 dcl2* double mutant were a gift of Dr. Martin Urban, Rothamsted Research, England. RNAi gene deletion mutants were generated in the *Fg* strain IFA65 (Jansen et al. 2005) hereafter termed IFA WT. They were generated by homolog recombination using the pPK2 binary vector. *Fg* RNAi genes were identified by blasting *Neurospora crassa* genes against the Fusarium genome sequence in the Broad institute data base. Disruption vectors were constructed by inserting two flanking fragments (∼1000 bp) upstream and downstream the corresponding genes in the pPK2 vector as follows: *RdRP1, AGO1, QDE3, QIP, AGO2, DCL1, RdRP2, RdRP3, RdRP4* and *DCL2* upstream flanking sequences were inserted in the plasmid between PacI-KpnI restriction sites, and the downstream flanking sequence were inserted between XbaI-HindIII restriction sites. Except *AGO2* downstream flanking sequence was inserted in XbaI restriction site (primers used in disruption plasmid construction are listed in Table S1. Disruption vectors were introduced into *Agrobacterium tumefaciens* (LBA440 and AGL1 strains) by electroporation. A single colony of Agrobacterium containing the pPK2 plasmid were grown in 10 ml YEB medium (Vervliet et al., 1975) containing the appropriate antibiotics (5 μg/ml tetracicllin + 25 μg/ml rifampicin + 50 μg/ml Kanamycin for LBA440, and 25 μg/ml carbenicillin + 25 μg/ml rifampicin + 50 μg/ml kanamycin for AGL1) and were incubated at 28°C till OD_600nm_ 0.7 was reached. T-DNA was mobilized in Agrobacterium with 200 μM acetosyringone, and Agrobacterium and fungal recipient IFA WT were co-cultivated on black filter paper (DP 551070, Albert LabScience, Hahnemühle, Dassel, Germany), respectively. Putative fungal mutants were selected on potato extract glucose (PEG) medium containing 150 μg/ml hygromycin + 150 μg/ml ticarcillin and grown for five days. For genotyping, genomic DNA of putative Fusarium mutants were extracted from mycelia.

### Genotyping of Fusarium mutants

*Fg* IFA mutants were confirmed by genotyping using primers located in hygromycin and corresponding gene flanking sequences (located after the cloned flanking sequence in the genome) (Table S2). Upon amplification the samples were sequenced. Additionally, mRNA expression levels of deleted vs. levels in IFA WT was measured by quantitative real time PCR (qRT-PCR) using primers pairs listed in (Table S3). The mRNA transcripts were measured using 1 x SYBR Green JumpStart Taq Ready Mix (Sigma-Aldrich, Germany) according to manufacturer’s instructions and assayed in 7500 Fast Real-Time PCR cycler (Applied Biosystems Inc, CA, USA) under the following thermal cycling conditions: initial activation step at 95°C for 5 min, 40 cycles (95°C for 30 s, 53°C for 30 s, and 72°C for 30 s). The Ct values were determined with the software in the qRT‐PCR instrument and the transcript levels of the genes was determined according to the 2^−ΔΔCt^ method (Livak and Schittgen, 2001).

### Colony morphology

The RNAi mutants were cultured on PEG (ROTH, Germany); starch agar (SA) and synthetic nutrient agar (SNA) (Leslie and Summerell, 2006). The cultures were incubated at 25°C in 12 h light/12 h dark (52 µmol m^−2^ s^−1^, Philips Master TL-D HF 16W/840). The growth was documented after 5 days. For growth in liquid cultures, agar blocks from two-week-old fungal cultures were incubated on liquid PEG medium for five days at room temperature (RT), light (2 µmol m^−2^ s^−1^) with shaking. Each mutant was grown in flask containing medium supplemented with hygromycin (100 μg/ml) and flask containing medium without hygromycin. Photos were taken to document the growth pattern after five days incubation.

### Production of fungal biomass

Fifty milligram mycelia (fresh mycelia from four-day-old fungal cultures grown on Aspergillus complete medium (CM) plates in the dark; Leslie and Summerell, 2006) were incubated in a 100 ml flask containing 20 ml of PEG medium incubated at RT with shaking under 12 h light (2 µmol m^−2^ s^−1^). Fungal mycelium was harvested after three days by filtration through filter paper (Munktell, AHLSTROM, Germany GMBH), washed with distilled water twice and dried at 75°C overnight. The dry weight was calculated by using the following formula: Dry weight = (weight of filter paper + mycelium) - (weight of filter paper).

### Conidiation assay

Production of conidia was done according to Yun et al., (2015) with slight modification. Four-day-old cultures of each mutant and IFA WT growing in CM agar plates in the dark at 25°C were used for fresh mycelia preparation. The mycelia were scraped from the plate surface using a sterile toothpick, then 50 mg mycelia were inoculated in a 100 ml flask containing 20 ml of SN medium. The flasks were incubated at RT for five days in light (2 µmol m^−2^ s^−1^) on a shaker (100 rpm). Subsequently, the conidia produced from each mutant and wt were counted using a hemocytometer (Fuchs Rosenthal, Superior Marienfeld, Germany).

### Viability test of conidia

Fourteen mL from the same cultures used in conidiation assay were centrifuged at 4,000 rpm for 10 min to precipitate conidia. The conidia were resuspended in 5 ml 2% sucrose water and incubated in dark for two days at 23°C. Germinated and non-germinated conidia were visualized and counted under an inverse microscope. Conidia germination rate was determined as percentage of germinated conidia of the total conidia number.

### Perithecia production and ascospore discharge assay

Fungi were grown on carrot agar prepared under bright fluorescent light at RT (18-24°C) for five days (Klittich and Leslie, 1988). Aerial mycelia were removed with a sterile tooth stick. To stimulate sexual reproduction and perithecia formation, one ml of 2.5% Tween 60 was applied to the plates with a sterile glass rod after scraping the mycelia (Cavinder et al., 2012). The plates were incubated under fluorescent light at RT for nine days. Subsequently, agar blocks (1.5 cm in diameter) were cut from the plates containing the mature perithecia using a cork borer. Agar blocks were sliced in half, placed on glass microscope slides, and incubated in boxes under high humidity for two days under 24 h light (52 µmol m^−2^ s^−1^ Philips Master TL-D HF 16W/840). During this time, ascospores discharged from the perithecia accumulated on the slide. For the quantification of discharged ascospores, slides were washed off by 2 ml of an aqueous Tween 20 (0.002%) solution and counted using a hemocytometer.

### Viability test of the discharged ascospores

Mycelia with mature perithecia (13 days after sexual induction) on carrot agar were incubated in a humid box at RT under light for four days according to Son et al., (2017). The discharged ascospores were washed from the plate cover using SN liquid medium and incubated in the dark for 24 h in a humid box. The germinated and non-germinated ascospores were visualized under an inverse microscope and counted.

### Pathogenicity assay on wheat ears

The susceptible wheat cultivar Apogee was used. Plants were grown in an environmentally controlled growth chamber (24°C, 16 h light, 180 μmol m^−2^ s^−1^ photon flux density, 60% rel. humidity) till anthesis. Point inoculations to the second single floret of each spike were performed at mid-anthesis with 5 μL of a 40,000 conidia/mL suspension amended with 0.002% v/v Tween 20 (Gosman et al., 2010). Control plants were treated with sterile Tween 20. For each *Fg* genotype, ten wheat heads were inoculated and incubated in plastic boxes misted with water to maintain high humidity for two days. Incubation continued at 22°C in 60% rel. humidity. Infected wheat heads were observed nine and 13 dpi and infection percentage was determined as the ratio of infected spikelets to the total spikelet number per ear.

### Thousand Grain Weight (TGW) of infected wheat kernels

A hundred kernels from two biological experiments with 10 wheat heads point-inoculated with IFA WT and mutants were counted and weighed. TGW was calculated in grams per 1000 kernels of cleaned wheat seeds.

### Quantification of fungal DNA in infected wheat kernels

Fungal genomic DNA in kernels was quantified using qPCR as described (Brandfass and Karlovsky, 2008). Dried grains were ground and DNA was extracted from 30 mg flour and dissolved in 50 µl of TE buffer. One µl of 50x diluted DNA was used as template for RT-PCR with primers amplifying a 280 bp fragment specific for *Fg*. The PCR mix consisted of reaction buffer (16 mM (NH_4_)_2_SO_4_, 67 mM Tris–HCl, 0.01% Tween-20, pH 8.8 at 25°C; 3 mM MgCl_2_, 0.3 μM of each primer, 0.2 mM of each dATP, dTTP, dCTP and dGTP (Bioline), 0.03 U/µl Taq DNA polymerase (Bioline, Luckenwalde, Germany) and 0.1x SYBR Green I solution (Invitrogen, Karlsruhe, Germany). The PCR was performed in CFX384 thermocycler (BioRad, Hercules, CA, USA) according to the following cycling condition: Initial denaturation 2 min at 95°C, 35 cycles with 30 s at 94°C, 30 s at 61°C, 30 s at 68°C, and final elongation for 5 min at 68°C. No matrix effects were detectable with 50-fold diluted DNA extracted from grains. Standards were prepared from pure *Fg* DNA in 3-fold dilution steps from 100 pg to 0.4 pg/well.

### Analysis of mycotoxins in infected wheat kernels

The content of mycotoxins in wheat kernels infected with *Fg* RNAi mutants and IFA WT was determined using high performance liquid chromatography coupled to tandem mass spectrometry (HPLC–MS/MS). Mycotoxins were extracted from ground grains with mixture containing 84% acetonitrile, 15% water and 1% acetic acid and the extracts were defatted with cyclohexane. Chromatographic separation was carried out on a C18 column eluted with a water/methanol gradient and the analytes were ionized by electrospray and detected by MS/MS in multiple reaction monitoring (MRM) mode essentially as described (Sulyok et al., 2006).

### Spray application of dsRNA on barley leaves

Second leaves of three-week-old barley cultivar Golden Promise were detached and transferred to square Petri plates containing 1% water-agar. dsRNA spray applications and leaf inoculation was done as described (Koch et al. 2016). For the TE-control, TE-buffer was diluted in 500 μl water corresponding to the amount used for dilution of the dsRNA. Typical RNA concentration after elution was 500 ng μl^−1^, representing a buffer concentration of 400 μM Tris-HCL and 40 μM EDTA in the final dilution. TE buffer were indistinguishable from treatments with control dsRNA generated from the GFP or GUS gene, respectively (Koch et al., 2016; Koch et al., 2018). Thus, we used TE buffer as control to save costs. Spraying of the leaves was carried out in the semi-systemic design (Koch et al. 2016), where the lower parts of the detached leaf segments were covered by a tinfoil to avoid direct contact of dsRNA with the leaf surface that was subsequently inoculated.

### Statistics and analysis

Data obtained from two or three repetitions were subjected to the Student’s *t* test in Microsoft office Excel 2010. Significance was determined as P≤0.05, 0.01 or 0.001 and indicated by *, ** or ***, respectively. Unless specified otherwise, data are presented as mean ± standard error or mean ± standard deviation of the mean. Sequence analysis was performed on the ApE plasmid editor free tool. Basic Local Alignment Search Tool (BLAST) NCBI BLAST (http://blast.ncbi.nlm.nih.gov/Blast.cgi) was used for sequences search and alignment.

## List of abbreviations

AGO: ARGONAUTE
CYP51: *Cytochrome P450 lanosterol C-14α-demethylase*
DCL: DICER-like
DON: deoxynivalenol
*Fg*: *Fusarium graminearum*
FHB: Fusarium head blight
HIGS: host-induced gene silencing
hpRNA: hairpin RNA
MSUD: meiotic silencing by unpaired DNA
NIV: nivalenol
PEG: potato extract glucose
QDE 2,3: Quelling defective 2,3
QIP: QDE-interacting protein
RdRp: RNA-dependent RNA polymerase
RISC: RNA-dependent silencing complex
RNAi: RNA interference
RPA: subunit of replication protein A
siRNA: small interfering RNA
SN: synthetic nutrient agar
ssDNA: single-stranded
TGW: thousand grain weight

## Declarations

### Ethics approval and consent to participate

Not applicable

### Consent for publication

Not applicable

### Availability of data and material

All data generated or analysed during this study are included in this published article [and its supplementary information files].

### Competing interests

The authors declare that they have no competing interests” in this section.

### Funding

This research was supported by the German Research Council (DFG) to K.-H. K and AK in the project GRK2355.

## Acknowledgements

We thank Mrs. E. Stein for excellent technical assistance, Dr. A. Rathgeb for mycotoxin analysis and Ms. C. Birkenstock for caring of the plants. We also thank Dr. Martin Urban, Rothamsted Research, England for providing the *Fg* strains PH1 and PH1 *dcl1 dcl2*.

## Supplement data

### Supplement figures

**Figure S1:**

**Schematic representation of the gene replacement strategy used for *F. graminearum* transformation.** Yellow box: the target gene that has to be replaced by KO; dark green box: selection marker gene, in this case the antibiotic resistance gene (*hygromycin B phosphotransferase* of *E. coli*, hph). Blue arrow: Homologous recombination sequences, typically ∼1 kb long; Black arrows: template area for primers binding used for transformants genotyping. PgpdA: Promoter region of the *Glyceraldehyde-3-phosphate dehydrogenase* gene of *Aspergillus nidulans*; TtrpC: termination region of the *Aspergillus nidulans trpC* gene.

**Fig. S2. Compromised expression of deleted RNAi genes in *F. graminearum* knockout (KO) mutants.** Expression of the targeted genes in respective Fusarium mutants. Transcript levels were analyzed by qRT-PCR from five-day-old PEG liquid cultures and transcript quantified by normalization to Fusarium *β-TUBULIN* (*FgTub*) or *ELONGATION FACTOR A* (*FgEF1a*) and comparison to IFA WT.

**Figure S3. Colony morphology and growth of RNAi KO mutants.** Fusarium mutants and wt IFA WT were grown for 5 days on solid (A) PDA (potato dextrose agar), (B) SN (synthetic nutrient), (C) CM (Aspergillus complete medium) and in liquid PEG medium without hygromycin. The mutants showed differences in pigmentation as follows: *Δago1, Δrdrp2, Δrdrp3* and *Δrdrp4* darker pigmentation; *Δdcl1, Δdcl2* and *Δrdrp1* reduced pigmentation compared to IFA WT.

**Fig. S4. Infection of wheat spikes with *F. graminearum* RNAi mutants and IFA WT.** Thousand grain weight (TGW) of infected wheat spikes. Mock control: Kernels treated with 0.002% Tween 20; mature kernels: completely mature Apogee kernels.

## Notes

#### Summary of Updates

A new table 2 has been uploaded for a Synopsis of the effects of mutations.

