## Supplementary figures and images for "Different components of the RNAi machinery are required for conidiation, ascosporogenesis, virulence, DON production and fungal inhibition by exogenous dsRNA in the Head Blight pathogen *Fusarium graminearum*"

### supplemental figures

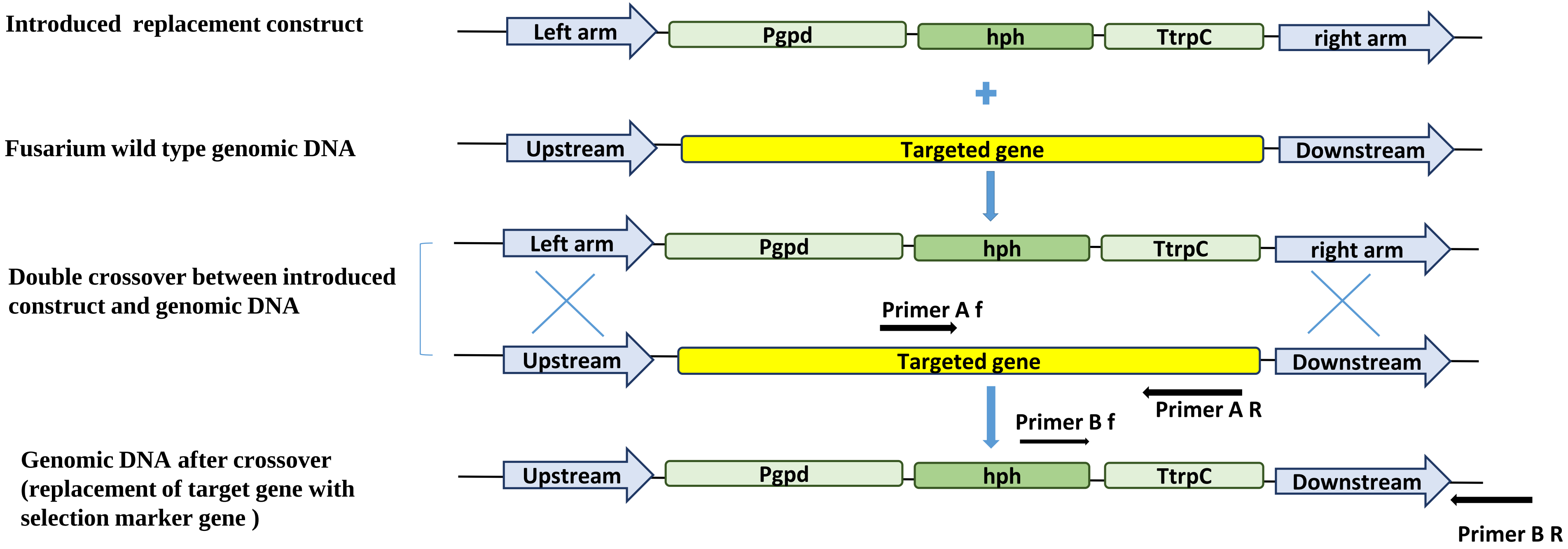

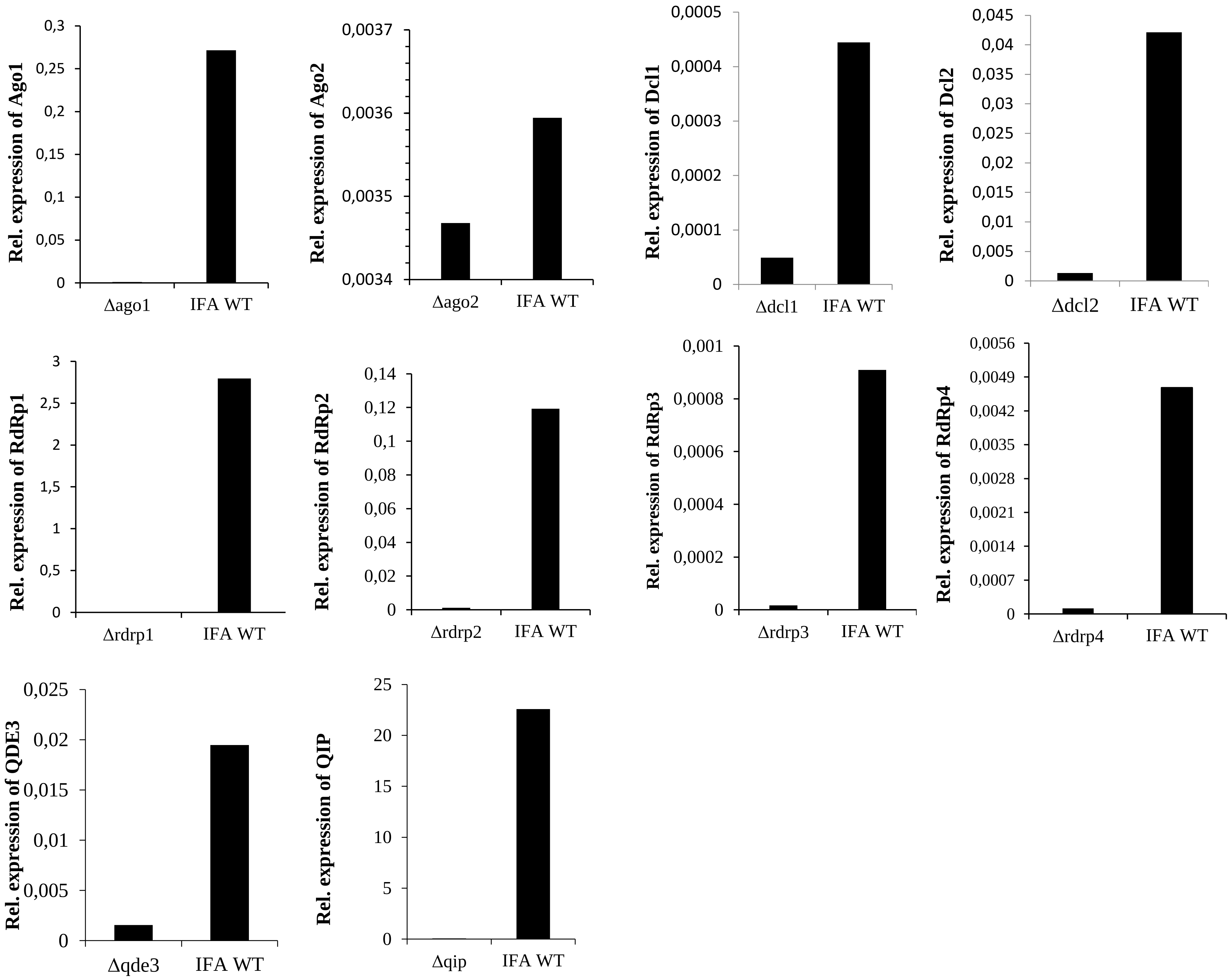

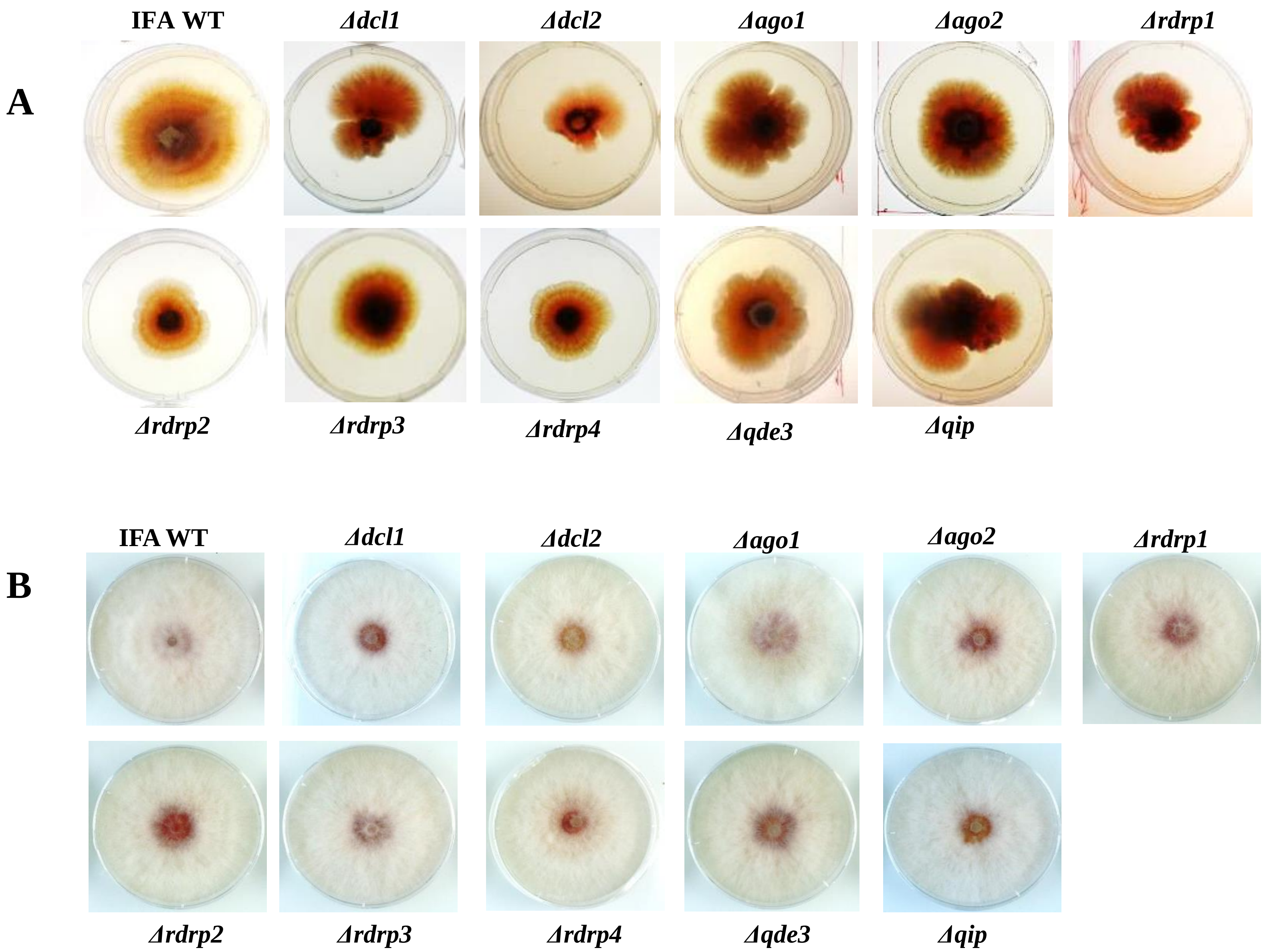

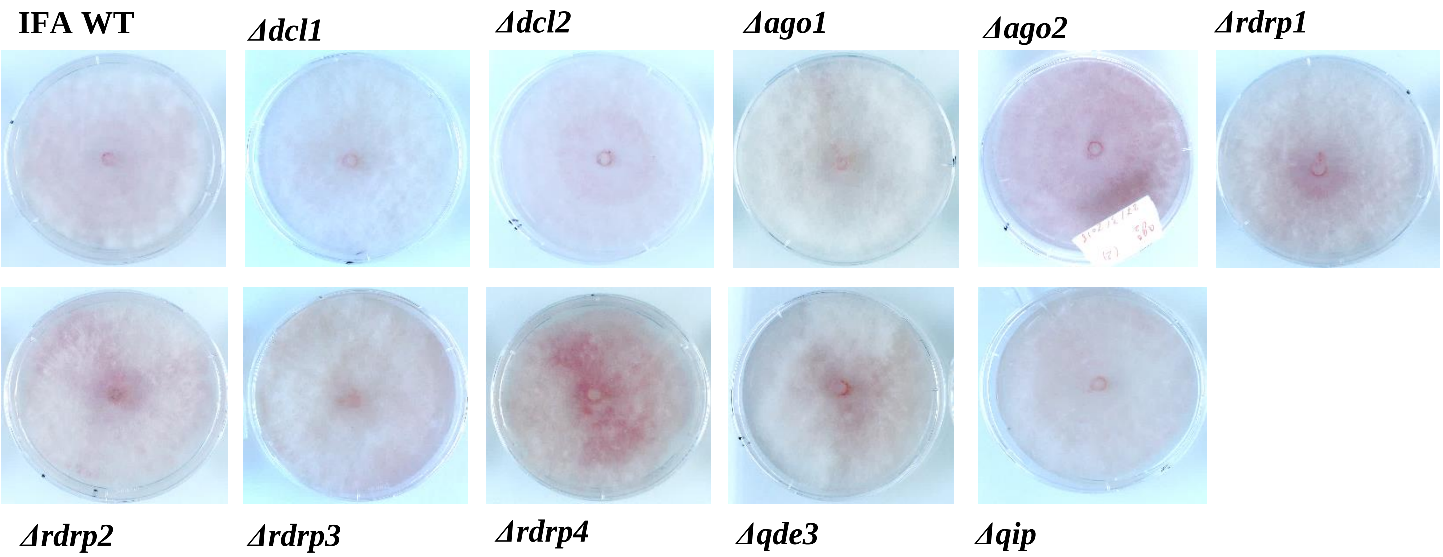

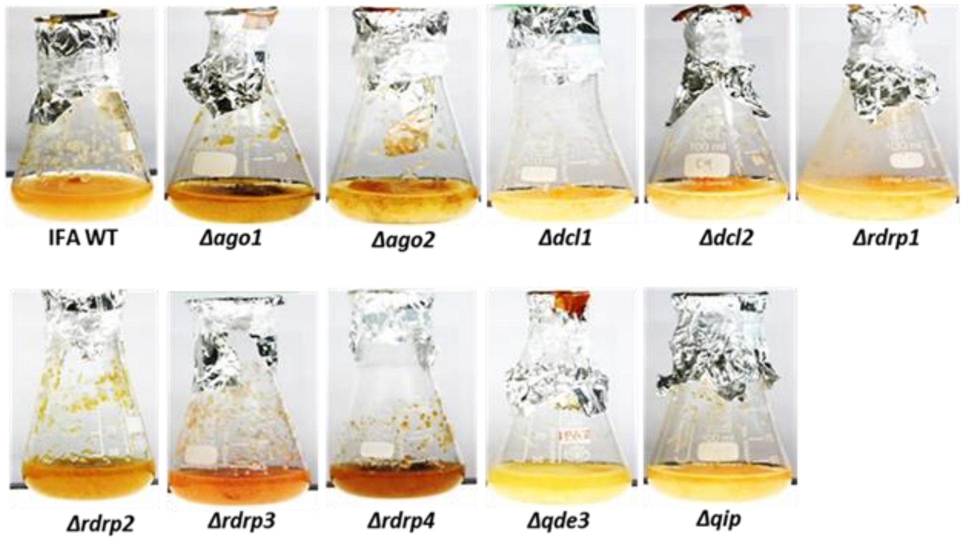

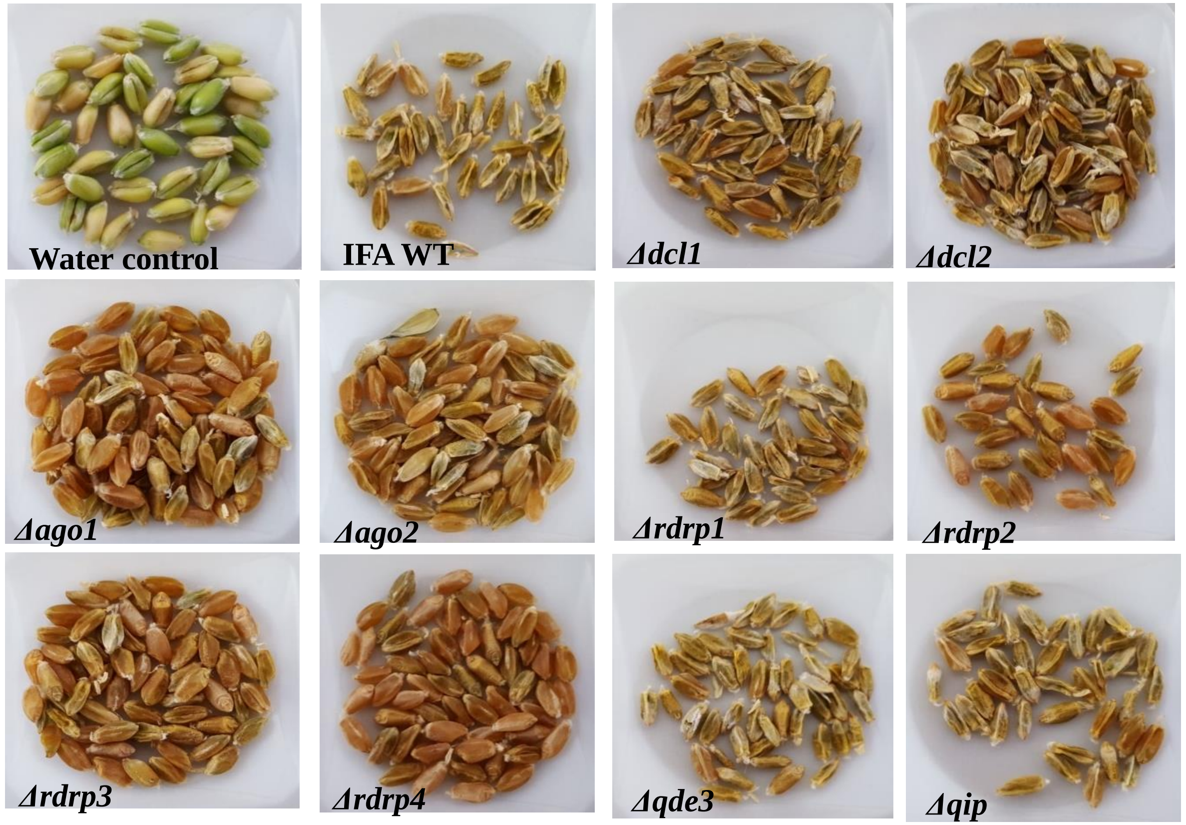

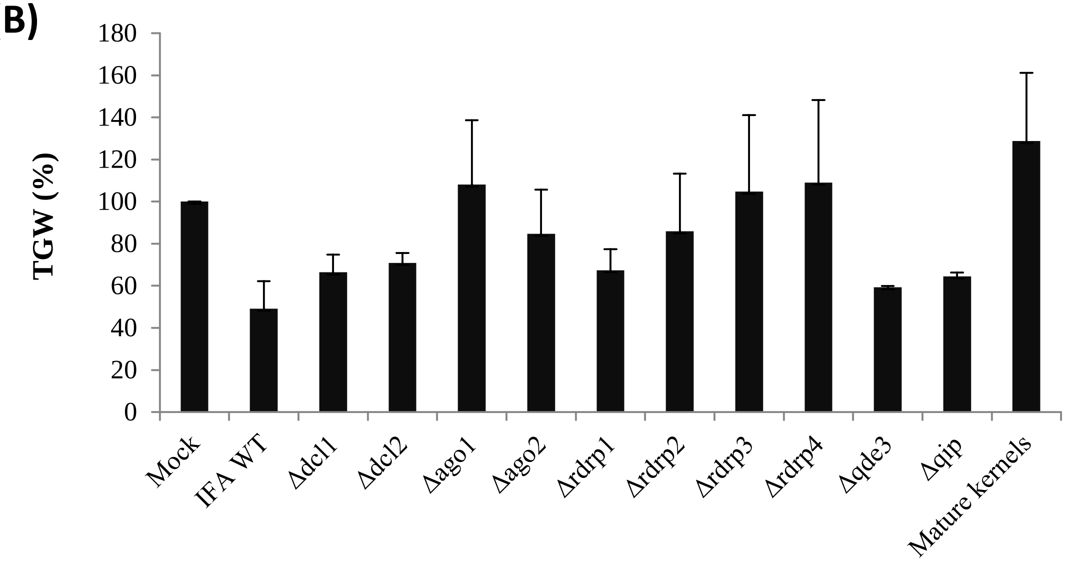
